# Tcf12 controls dynamic calvarial bone growth and motor learning in mice

**DOI:** 10.1101/2024.01.09.574781

**Authors:** Takahiko Yamada, Jesse Anderson-Ramirez, Lu Gao, Peng Chen, Mingyi Zhang, Tingwei Guo, Jifan Feng, Thach-Vu Ho, Ivetta Vorobyova, Naomi Santa Maria, Russell Jacobs, Jian-Fu Chen, Yang Chai

## Abstract

Heterozygous loss-of-function mutations of *TCF12* and *TWIST1* can each cause craniosynostosis and neurodevelopmental delay in humans. Twist1-Tcf12 interaction plays an important role in regulating suture development. Although the molecular and cellular mechanisms underlying craniosynostosis and neurocognitive dysfunctions in *Twist1*^+/-^ mice have been studied, less information on the role of *Tcf12* in these defects is available. To investigate the functional mechanism of Tcf12 in regulating skull and brain development, we analyzed the skull shape of *Wnt1-Cre;Mesp1-Cre;Tcf12^fl/fl^* mice and found that, despite mild coronal synostosis, their skull shape appears to be similar to that of controls. We also found evidence of impaired motor learning ability in *Tcf12* mutant mice. Furthermore, loss of Tcf12 in neural crest lineage leads to upregulated *Runx2* expression in the calvarial mesenchyme and posterior expansion of the frontal bone in *Wnt1-Cre;Tcf12^fl/fl^* mice. Mechanistically, we show that *Lmx1b* is a direct downstream target of Tcf12 for the regulation of osteogenic differentiation in the calvarial mesenchyme during embryonic development. Importantly, overexpression of *Lmx1b* inhibits osteogenic differentiation in the calvarial mesenchyme of *Wnt1-Cre;Tcf12^fl/fl^* mice, indicating Tcf12’s regulation of *Lmx1b* expression is crucial for controlling osteogenesis during calvarial bone development. Our study suggests that Tcf12 expression in the brain is crucial for motor learning. Moreover, this study establishes a new molecular mechanism underlying regulation of calvarial bone formation.

**Author Summary:** Craniosynostosis is characterized by premature fusion of cranial sutures and associated with abnormal skull growth, delayed brain development, and often impaired brain functions. Loss-of-function mutation of *TCF12* can cause coronal synostosis and neurodevelopmental delay in humans. In developing mouse sutures, *Tcf12* is essential for maintaining the boundary between sutural and osteogenic cells. However, roles of *Tcf12* in skull formation and brain development have not been fully investigated. In this study, we show that loss of *Tcf12* leads to brain abnormalities even in the absence of coronal synostosis and that frontal bone expansion results from upregulated osteogenic differentiation in the calvarial mesenchyme in mice. Furthermore, we identify *Lmx1b* as a downstream target of Tcf12 for the regulation of osteogenic differentiation in the calvarial mesenchyme during frontal bone development. Our findings highlight the role of Tcf12 in the development of calvarial bones and provide new insight into molecular mechanisms for regulation of calvarial bone formation.

## Introduction

Cranial sutures–which include the inter-frontal (metopic), coronal, lambdoid, and sagittal sutures–are fibrous joints that serve to connect the bones of the skull vault, coordinate growth and development of the skull and the brain, allow a small amount of movement, and absorb shock [1–3]. Craniosynostosis, the premature fusion of at least one cranial suture, is a common congenital defect found in 1 in 2,000-2,500 live births in humans. It is associated with abnormal skull growth, increased intracranial pressure (ICP), and abnormal or delayed brain development that may result in cognitive impairment [2–6].

In Saethre-Chotzen syndrome, the coronal suture joining the frontal and parietal bones is fused. This syndrome can be caused by heterozygous loss-of-function mutations in one of two basic helix-loop-helix (bHLH) transcription factor genes, *TWIST1* and *TCF12* [7–9]. Saethre-Chotzen syndrome displays autosomal dominant inheritance and occurs in 1/50,000 live births. Alongside coronal suture fusion, patients may exhibit abnormalities of the digits, facial dysmorphology, and/or intellectual disability [2]. Twist1 has the ability to form homodimers but can also make heterodimers with other bHLH factors, including Tcf12, a member of the E-protein subfamily [10]. It has been reported that a higher Twist1/Twist1 to Twist1/E-protein dimerization ratio is associated with an expansion of the osteogenic fronts and suture closure in mice [11]. The specific role that *Tcf12* plays in skull development remains unclear, as does whether *Twist1* and *Tcf12* have overlapping or distinct regulatory functions during suture and calvarial development processes.

Heterozygous loss of *Twist1* in mice produces coronal suture fusion similar to that seen in human Saethre-Chotzen syndrome patients [8], although unlike in humans, heterozygous deletion of *Tcf12* does not cause suture defects in mice [9]. Recently, it was shown that tissue-specific homozygous loss of *Tcf12* results in several of the same abnormal calvarial phenotypes as the heterozygous loss of *Twist1*, including coronal suture fusion, unusually rapid frontal and parietal bone growth, and a perturbed boundary between the suture and the neighboring osteogenic cell population [12]. A previous study established that *Twist1* heterozygous mice with coronal suture fusion exhibit neurocognitive behavioral abnormalities [13]. However, whether loss of *Tcf12* leads to behavioral abnormalities has not yet been studied.

During embryonic development, the calvarial bones develop from the cranial mesenchyme layer that covers the brain, which contains cells from the neural crest and the mesoderm [1,14]. Neural crest-derived cells (NCCs) give rise to the frontal bone, inter-frontal suture, and sagittal suture, whereas mesoderm-derived cells give rise to the parietal bone and coronal suture mesenchyme [15,16]. Calvarial bone development in mice begins on embryonic day (E)12, with the emergence of the frontal and parietal bone rudiments in the basolateral cranial mesenchyme just above the eye, termed the supraorbital mesenchyme [1,14,17,18]. After osteogenesis begins in the supraorbital mesenchyme, the frontal and parietal bones grow toward the apex over the following days. No ossification centers arise from the early migrating mesenchyme, which is apical to the supraorbital mesenchyme [19]. Previous reports have shown that *Lmx1b* (LIM homeobox transcription factor 1 beta) is specifically expressed in the early migrating mesenchyme, where it inhibits osteogenesis and plays a crucial role in the apical-basal patterning of the calvaria [20,21]. Deletion of *Lmx1b* in the cranial mesenchyme leads to heterotopic ossification of the calvaria and the fusion of multiple sutures [20].

In this study, we investigated the functional mechanism of Tcf12 in regulating brain and skull development and discovered that loss of *Tcf12* leads to cerebellum development defect and abnormalities in motor function, suggesting that Tcf12 plays a key role in regulating cerebellum development. Additionally, we investigated the striking phenotype seen in the frontal bones of mice with deletion of *Tcf12* in neural crest cells to obtain insights into *Tcf12*’s functional mechanism in regulating osteogenesis through *Lmx1b* expression during calvarial bone development.

## Results

### Loss of *Tcf12* in NCCs and/or mesoderm-derived cells does not alter skull shape

We have previously shown that *Twist1^+/-^* mice with bilateral coronal suture fusion exhibit skull shape deformities and neurocognitive abnormalities [13]. Another study reported that homozygous loss of *Tcf12* in both cells derived from the neural crest and from the mesoderm (*Wnt1-Cre;Mesp1-Cre;Tcf12^fl/fl^*) causes coronal suture fusion [12]. To investigate whether coronal suture fusion can lead to skull shape deformities and neurocognitive abnormalities in *Wnt1-Cre;Mesp1-Cre;Tcf12^fl/fl^* mice, as reported in humans [9], we performed micro-computed tomography (µCT) analysis and behavioral tests of adult mice with deletion of *Tcf12* in NCCs (*Wnt1-Cre;Tcf12^fl/fl^*), mesoderm-derived cells (*Mesp1-Cre;Tcf12^fl/fl^*), or both (*Wnt1-Cre;Mesp1-Cre;Tcf12^fl/fl^*). We pursued this conditional knockout strategy since *Tcf12^+/-^* mice show no abnormal calvarial phenotype and *Tcf12^-/-^* mice do not survive more than 2 weeks after birth [22].

We first confirmed *Tcf12* deletion efficiency in adult mutant mice (Fig 1). Using RNAscope of 1-month-old (P1M) control mouse skulls, we observed expression of *Tcf12* in the coronal suture mesenchyme (Fig 1A). Since the coronal suture mesenchyme is derived from the mesoderm, we confirmed that *Tcf12* expression was efficiently deleted in the coronal suture mesenchyme of *Mesp1-Cre;Tcf12^fl/fl^* and *Wnt1-Cre;Mesp1-Cre;Tcf12^fl/fl^* mice (Fig 1C and 1D). Next, we carried out detailed skull shape analysis using µCT images of P1M mice. We compared the skull shapes of controls, *Wnt1-Cre;Tcf12^fl/fl^*, *Mesp1-Cre;Tcf12^fl/fl^*, and *Wnt1-Cre;Mesp1-Cre;Tcf12^fl/fl^* mice. Anatomical landmarks (S1 Fig) were used to analyze the shape of the calvaria viewed from the top (Fig 2A-2D) and from the side (Fig 2F-2I) in each group. Wireframes drawn from the skull shapes of each group were similar from both the top and lateral perspectives (Fig 2E and 2J). Principal component analysis (PCA) revealed that all groups, including the mice with coronal suture fusion, were similar to each other in both top and lateral shape (Fig 2K and 2L). µCT images of each group confirmed the coronal suture fusion in *Wnt1-Cre;Mesp1-Cre;Tcf12^fl/fl^* mice (Fig 2D; arrowhead) and that the position of the Bregma (the junction of the sagittal and coronal sutures) shifted posteriorly in *Wnt1-Cre;Tcf12^fl/fl^* and *Wnt1-Cre;Mesp1-Cre;Tcf12^fl/fl^* mice (Fig 2B and 2D; arrows). The ratio of the sagittal suture length to the metopic suture length in *Wnt1-Cre;Tcf12^fl/fl^* and *Wnt1-Cre;Mesp1-Cre;Tcf12^fl/fl^* mice was less than in the control (Fig 2M and 2N), which is consistent with a previous report [12]. These results indicated that deletion of *Tcf12* in neural crest- and mesoderm-derived cells led to partial coronal suture fusion, which was not enough to cause skull shape deformities and represented a less severe phenotype than what is typically seen in *Twist1^+/-^* mice. Also, our analysis suggested that the frontal bone was elongated toward the posterior direction following *Tcf12* deletion in NCCs.

**Fig 1.**
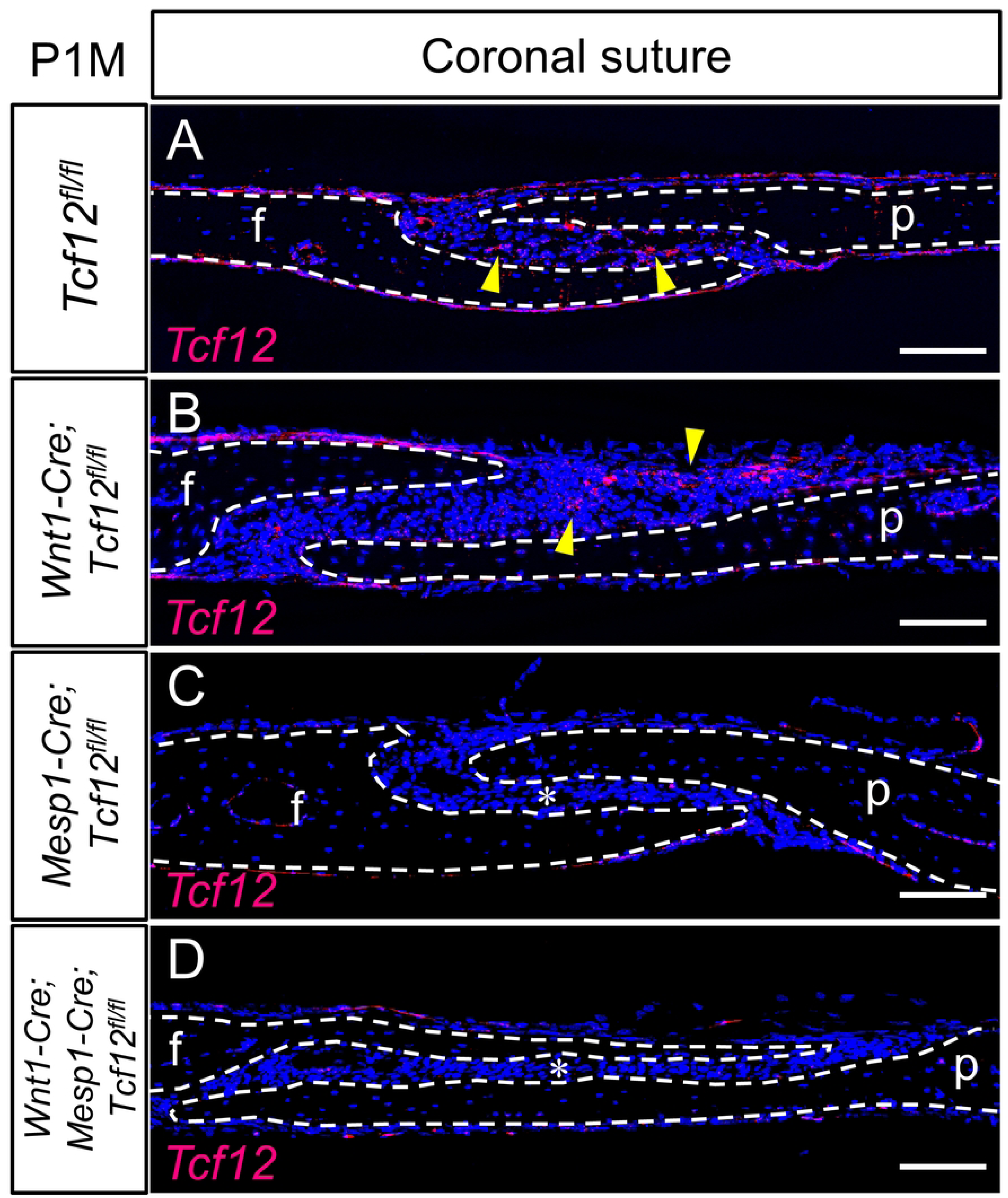
Tissue-specific deletion of *Tcf12* in the coronal suture of adult mice. (A-D) *Tcf12* expression pattern in the coronal suture at P1M in *Tcf12^fl/fl^* (A), *Wnt1-Cre;Tcf12^fl/fl^* (B), *Mesp1-Cre;Tcf12^fl/fl^* (C), and *Wnt1-Cre;Mesp1-Cre;Tcf12^fl/fl^* (D) mice. Frontal bones and parietal bones are outlined by white dashed lines in A-D. Yellow arrowheads in A and B point to the *Tcf12* signals. Asterisks in C and D indicate the deleted expression of *Tcf12* in the coronal suture mesenchyme. f, frontal bone; p, parietal bone. Scale bar = 200 μm.

**Fig 2.**
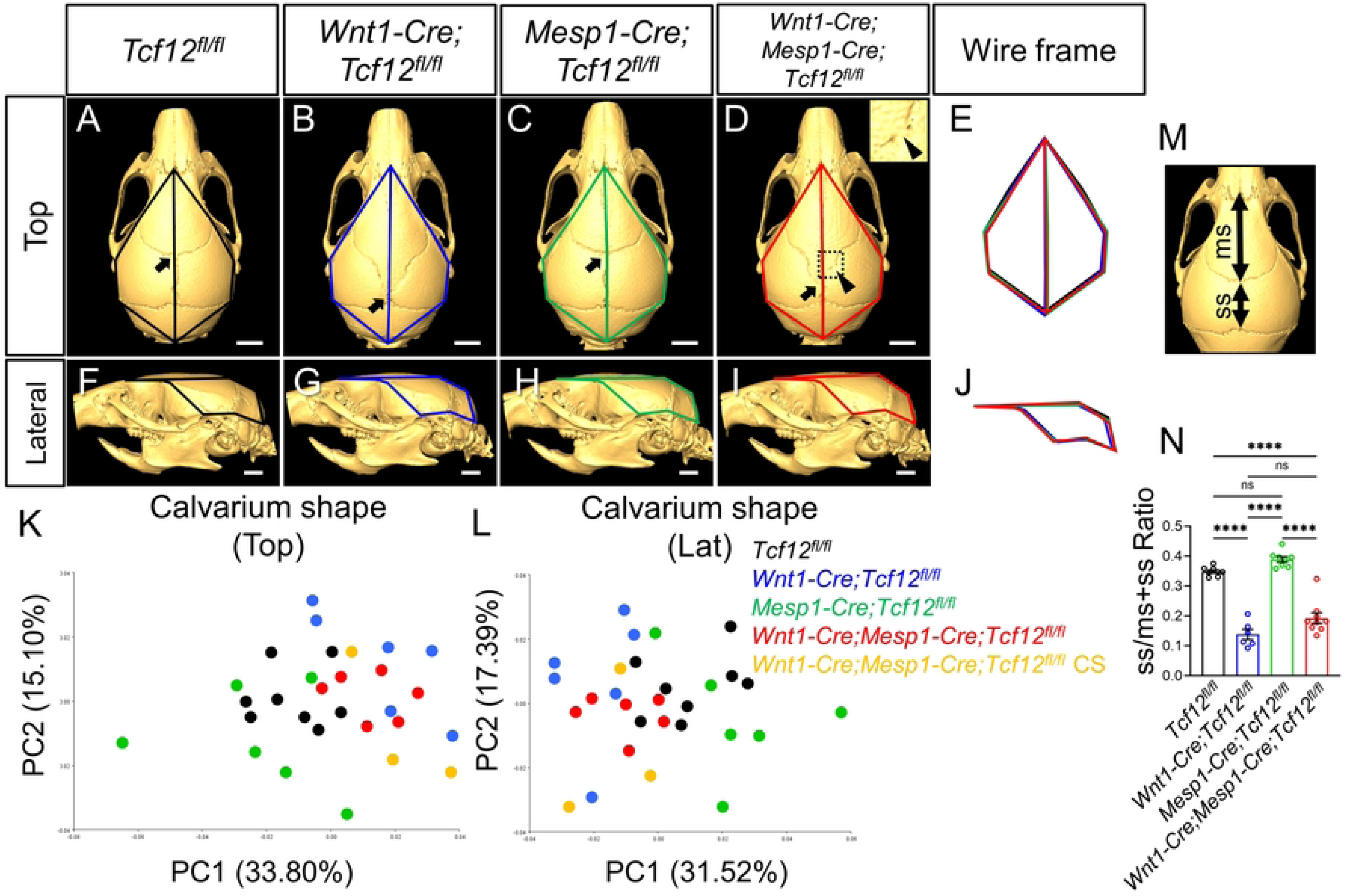
Loss of *Tcf12* in NCCs and mesoderm-derived cells causes coronal synostosis without affecting skull shape. (A-J) 3D μCT images with wireframes of top (A-D) and lateral (F-I) views of the head at P1M from *Tcf12^fl/fl^* (A, F), *Wnt1-Cre;Tcf12^fl/fl^* (B, G), *Mesp1-Cre;Tcf12^fl/fl^* (C, H), and *Wnt1-Cre;Mesp1-Cre;Tcf12^fl/fl^* (D, I) mice. The panel at top right in D shows higher magnification of the white dotted rectangle area. Wireframes in black, blue, green, and red were drawn according to landmarks for *Tcf12^fl/fl^*, *Wnt1-Cre;Tcf12^fl/fl^*, *Mesp1-Cre;Tcf12^fl/fl^*, and *Wnt1-Cre;Mesp1-Cre;Tcf12^fl/fl^* mice, respectively. Wireframe comparison representing the shape differences between 4 groups from top (E) and lateral (J) views. Black arrows in A-D point to the Bregma. Black arrowhead in D points to the fused coronal suture area. (K, L) Total variation between *Tcf12^fl/fl^* (black, n=8), *Wnt1-Cre;Tcf12^fl/fl^* (blue, n=6), *Mesp1-Cre;Tcf12^fl/fl^* (green, n=6), *Wnt1-Cre;Mesp1-Cre;Tcf12^fl/fl^* without coronal suture fusion (red, n=6), and *Wnt1-Cre;Mesp1-Cre;Tcf12^fl/fl^* with coronal suture fusion (orange, n=3) mice at P1M was determined by PCA for top (K) and lateral (L) views. (M, N) Comparison of the ratio of the length of sagittal suture (ss) to the metopic suture (ms) at P1M. Schematic diagram of the method for calculating ss/ms+ss ratio (M). Quantification of the ratio of the sagittal suture to the metopic suture in *Tcf12^fl/fl^* (black, n=8), *Wnt1-Cre;Tcf12^fl/fl^* (blue, n=6), *Mesp1-Cre;Tcf12^fl/fl^* (green, n=6), and *Wnt1-Cre;Mesp1-Cre;Tcf12^fl/fl^* (red, n=9) mice (N). Statistical differences were assessed with one-way ANOVA; **** = *P*-value <.0001; ns, not significant. Error bars represent s.e.m. Scale bar = 2 mm (A-D, F-I).

### Loss of *Tcf12* in *Wnt1-*expressing cells leads to small cerebellum and impaired motor learning ability in mice

To study whether deletion of *Tcf12* can cause similar cognitive defects to those seen in *Twist1^+/-^* mice, we conducted standard cognitive behavioral tests to assess object memory, short-term spatial memory, anxiety, and social cognition. We found no significant differences between groups on this battery of tests (S2A-2E Fig). We then examined motor learning ability using a rotarod test (Fig 3A). Control mice were able to spend increasingly more time on the accelerating rotarod before falling off over four consecutive days, indicating active motor learning (Fig 3B; black). In contrast, *Wnt1-Cre;Tcf12^fl/fl^* and *Wnt1-Cre;Mesp1-Cre;Tcf12^fl/fl^* mice did not exhibit obvious improvement (Fig 3B; blue and red), suggesting motor learning impairment. To investigate why this might be the case, we measured their brain volume using magnetic resonance imaging (MRI). Whole-brain volume, especially cerebellar volume, was reduced in *Wnt1-Cre;Tcf12^fl/fl^* and *Wnt1-Cre;Mesp1-Cre;Tcf12^fl/fl^* mice, while other assessed regions including as the cortex and hippocampus were not reduced in size (Figs 3C-3H, S2F and S2G). To determine the cellular basis of these brain volume changes, we examined cell proliferation activity in each region of the brain at E18.5. Immunohistochemistry (IHC) staining showed that Ki67, which is a proliferation marker, was significantly reduced in the cerebellum of *Wnt1-Cre;Tcf12^fl/fl^* mice while other regions were unaffected (Figs 3I-3K and S2H-2K). To confirm the *Tcf12* expression pattern during brain development, we also carried out *in situ* hybridization (ISH) at E18.5, revealing *Tcf12* expression in the cortex and hippocampus but not in the cerebellum (S2L and 2M Fig). These results suggested that deletion of *Tcf12* in the progeny of *Wnt1*-positive cells reduced proliferation in the cerebellum during brain development and led to reduction in the size of the cerebellum, causing impaired motor learning ability in *Wnt1-Cre;Tcf12^fl/fl^* and *Wnt1-Cre;Mesp1-Cre;Tcf12^fl/fl^* mice. Our findings also indicated that deletion of *Tcf12* can lead to impaired brain function independent of suture fusion.

**Fig 3.**
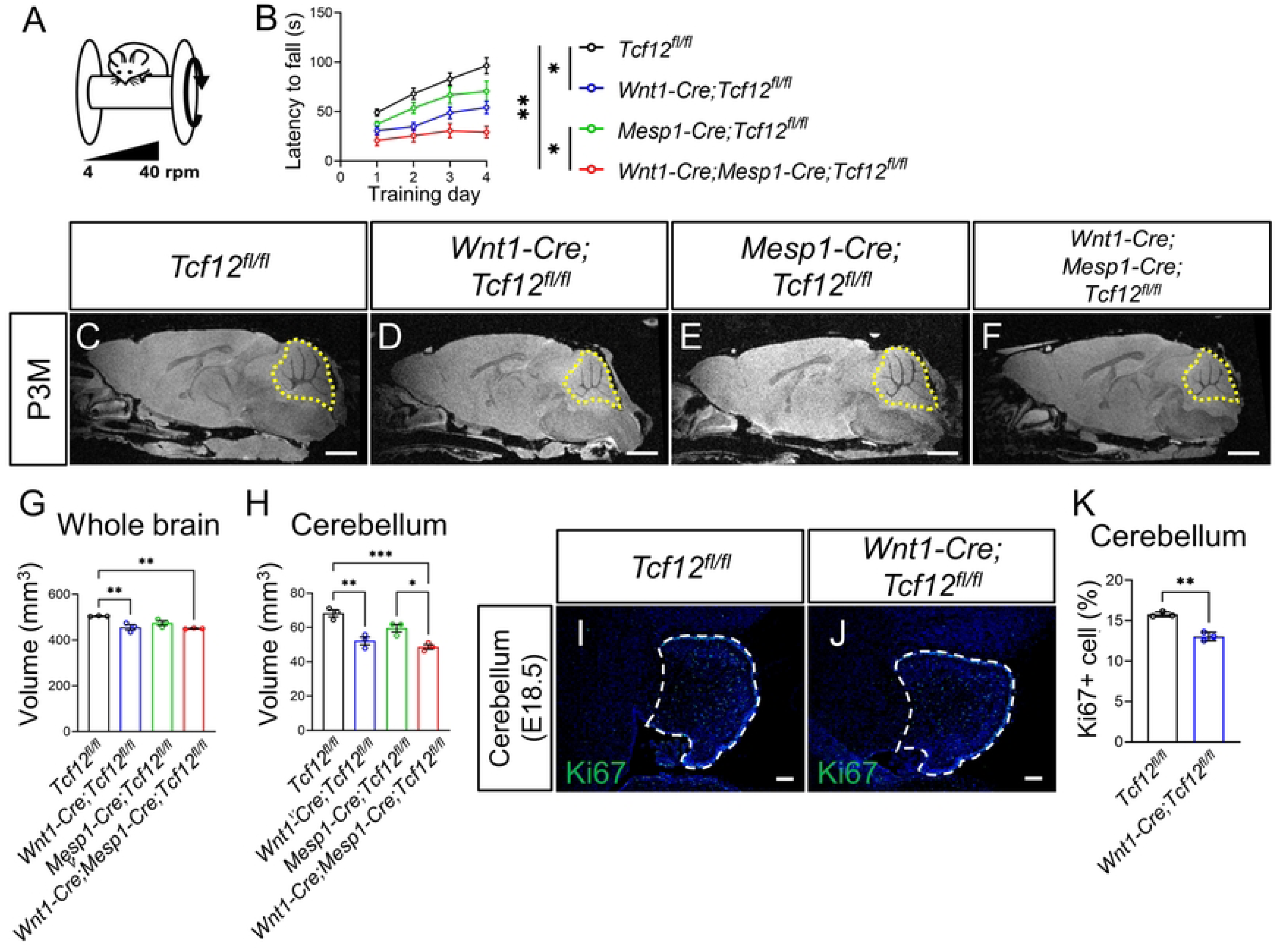
Loss of *Tcf12* in *Wnt1-*expressing cells leads to small cerebellum and impaired motor learning ability. (A, B) Schematic diagram of the rotarod test (A). Rotarod performance scored as time (seconds) on the rotarod for *Tcf12^fl/fl^* (black), *Wnt1-Cre;Tcf12^fl/fl^* (blue), *Mesp1-Cre;Tcf12^fl/fl^* (green), and *Wnt1-Cre;Mesp1-Cre;Tcf12^fl/fl^* (red) mice at P2M-P3M (B). Each group, n=20 including 10 males and 10 females. (C-H) MRI images of *Tcf12^fl/fl^* (C), *Wnt1-Cre;Tcf12^fl/fl^* (D), *Mesp1-Cre;Tcf12^fl/fl^* (E), and *Wnt1-Cre;Mesp1-Cre;Tcf12^fl/fl^* (F) mouse brains at P3M. Cerebellums are outlined by yellow dotted lines. Quantification of the volume of the whole brain (G) and the cerebellum (H) in *Tcf12^fl/fl^* (n=3), *Wnt1-Cre;Tcf12^fl/fl^* (n=3), *Mesp1-Cre;Tcf12^fl/fl^* (n=3), and *Wnt1-Cre;Mesp1-Cre;Tcf12^fl/fl^* (n=3) mice at P3M. (I-K) Ki67 (green) immunofluorescence on the cerebellum in *Tcf12^fl/fl^* (I) and *Wnt1-Cre;Tcf12^fl/fl^* (J) mouse embryos at E18.5. Cerebellums are outlined by white dashed lines. Quantification of the Ki67+ cells in the cerebellum of *Tcf12^fl/fl^* (n=3) and *Wnt1-Cre;Tcf12^fl/fl^* (n=3) mouse embryos at E18.5 (K). Ki67+ cell (%) is the percentage of Ki67 positive cells out of all cells in cerebellum outlined by white dashed lines in I and J. Statistical differences were assessed with one-way ANOVA (B, G, H) or unpaired two-tailed Student’s t-test (K); *** = *P*-value <.001; ** = <.01; * = <.05. Error bars represent s.e.m. Scale bars = 2 mm (C-F), 100 μm (I, J).

### Loss of *Tcf12* leads to frontal bone elongation and ectopic bone formation in *Wnt1-Cre;Tcf12^fl/fl^* mice

Loss of *Tcf12* in NCCs led to posterior expansion of the frontal bone paired with correspondingly decreased expansion of the parietal bone. To investigate the molecular and cellular mechanism responsible for this defect, we first compared frontal bone and coronal suture morphology in control and *Wnt1-Cre;Tcf12^fl/fl^* mice at newborn (P0), P1M, and P3M with bone staining and µCT (Fig 4A, B, C, D, H and I). Bone staining of P0 mice also showed ectopic bone formation in the medial part of the coronal suture area (Fig 4A and 4B), which is consistent with a previous report [12]. Frontal bone elongation toward the posterior of the skull was seen in the medial part of the coronal suture area (Fig 4D and 4I; asterisks) while areas of ectopic bone, which were not fused to any other calvarial bones, formed in the same area (Fig 4D and 4I; red arrowheads) at P1M. Interestingly, the overlap between the frontal bone and parietal bone in the coronal suture was flipped in the area of where this frontal bone elongation and ectopic bone formation were seen in *Wnt1-Cre;Tcf12^fl/fl^* mice (Fig 4F, 4K, 4G and 4L). These findings were confirmed in µCT sections and histological staining, which showed that the expanded frontal bones and ectopic bones did not fuse to the parietal bones. These results indicated that an altered osteogenic center in the posterior part of the frontal bone caused the ectopic bone formation, and then the ectopic bone fused with the frontal bone at a later stage to create what we later observed as frontal bone elongation.

**Fig 4.**
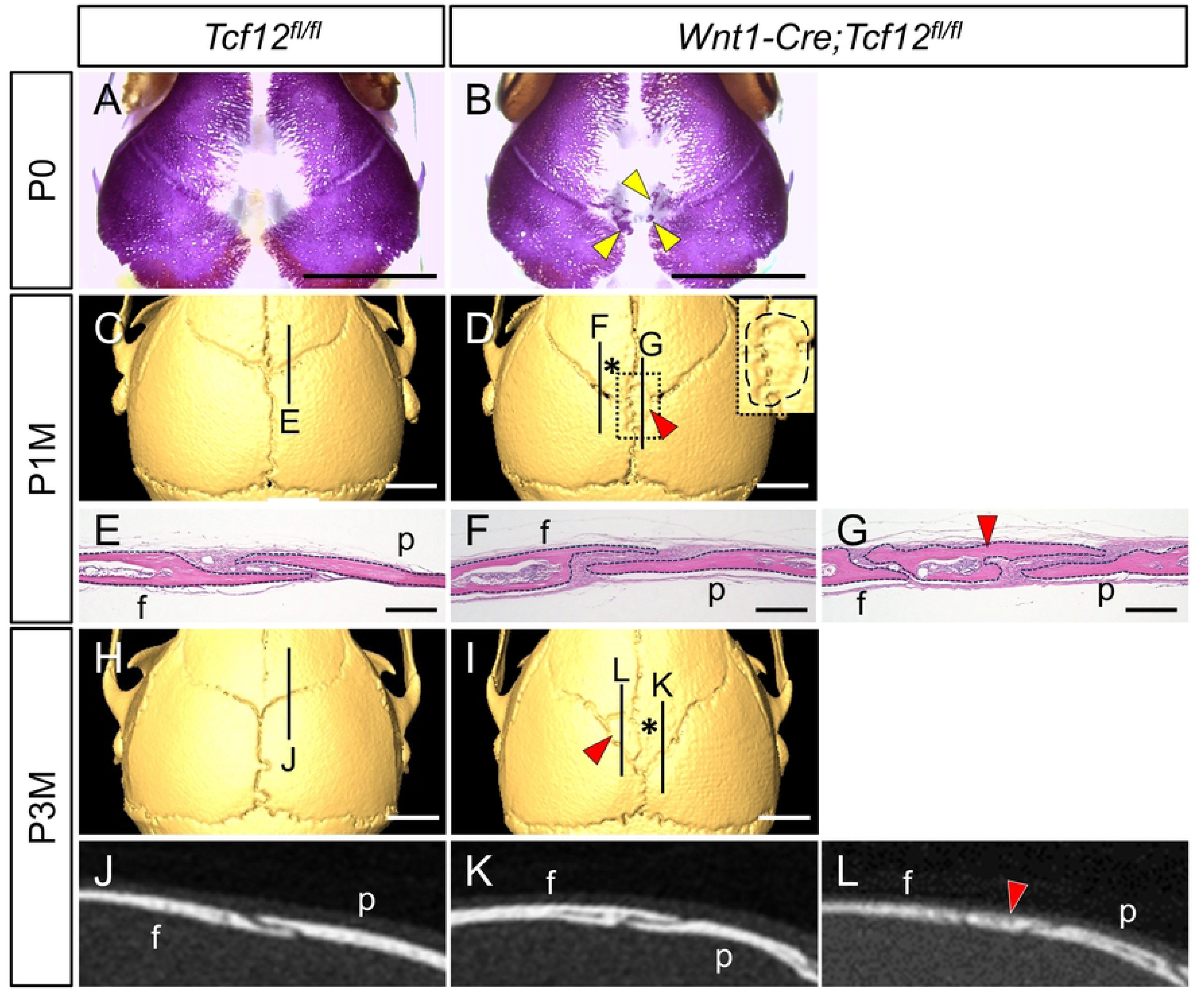
Loss of *Tcf12* in NCCs leads to frontal bone elongation and ectopic bones. (A, B) Alizarin Red S staining of P0 skulls from *Tcf12^fl/fl^* (A) and *Wnt1-Cre;Tcf12^fl/fl^* (B) mice. Yellow arrowheads point to the ectopic bones in B. (C-G) µCT images of the coronal suture at P1M in *Tcf12^fl/fl^* and *Wnt1-Cre;Tcf12^fl/fl^* mice (C, D). The panel at top right shows higher magnification of the white dotted rectangle area. The ectopic bone is circled by a black dashed line in E-G. (H-L) Hematoxylin and eosin staining of the coronal suture in *Tcf12^fl/fl^* (E) and *Wnt1-Cre;Tcf12^fl/fl^* (F, G) mice at P1M. Frontal, parietal, and ectopic bones are outlined by black dashed lines in E-G. (H-L) Micro-CT images of the coronal suture at P3M in *Tcf12^fl/fl^* and *Wnt1-Cre;Tcf12^fl/fl^* mice showing 3D reconstructions (H, I) and slices (J-L). Overlap of the frontal and parietal bones in the medial part of the coronal suture is inverted in F and K. Asterisks in D and I indicate the expanded area of the frontal bone. Red arrowheads point to the ectopic bones in D, G, I, and L. f, frontal bone; p, parietal bone. Scale bars = 2 mm (A-D, H, I), 100 μm (E-G).

### Osteogenic differentiation is upregulated in the medial mesenchyme of the frontal bone in *Wnt1-Cre;Tcf12^fl/fl^* mice

Since the ectopic bone was already formed at P0 (Fig 4B), we next analyzed cellular changes in the ectopic bone formation area at E15.5. In the meantime, we confirmed that *Tcf12* was expressed in the calvarial mesenchyme including the dura of control mice at E15.5 (S3B Fig), and that *Tcf12* expression was efficiently deleted in the calvarial mesenchyme and the dura of *Wnt1-Cre;Tcf12^fl/fl^* mice (S3C Fig). To investigate the cellular basis of the ectopic bone formation in the mutant mice, we examined osteogenic differentiation and cell proliferation in the posterior portions of the inter-frontal mesenchyme area and coronal suture area at E15.5 (Fig 5C-5F). IHC staining showed that the number of Runx2+ cells was increased in the posterior region of the inter-frontal mesenchyme and the most posterior edge of the frontal bone in *Wnt1-Cre;Tcf12^fl/fl^* mice (Fig 5G and 5J). In addition, the number of Runx2+ Ki67+ cells was also increased in the posterior region of the inter-frontal mesenchyme of *Wnt1-Cre;Tcf12^fl/fl^* mice, but not in the frontal bone edge (Fig 5H and 5K). The number of Runx2-Ki67+ cells was unaffected (Fig 5I and 5L). Osteogenic differentiation and cell proliferation in the coronal suture area were not affected in *Wnt1-Cre;Tcf12^fl/fl^* mice at E15.5 (S4 Fig). These results suggested that the deletion of *Tcf12* in the posterior portions of the inter-frontal mesenchyme and frontal bone led to upregulated osteogenic differentiation and proliferation of early osteoblasts in the mesenchyme and forming edge of the posterior frontal bones, causing an ectopic osteogenic center in the calvarial mesenchyme.

**Fig 5.**
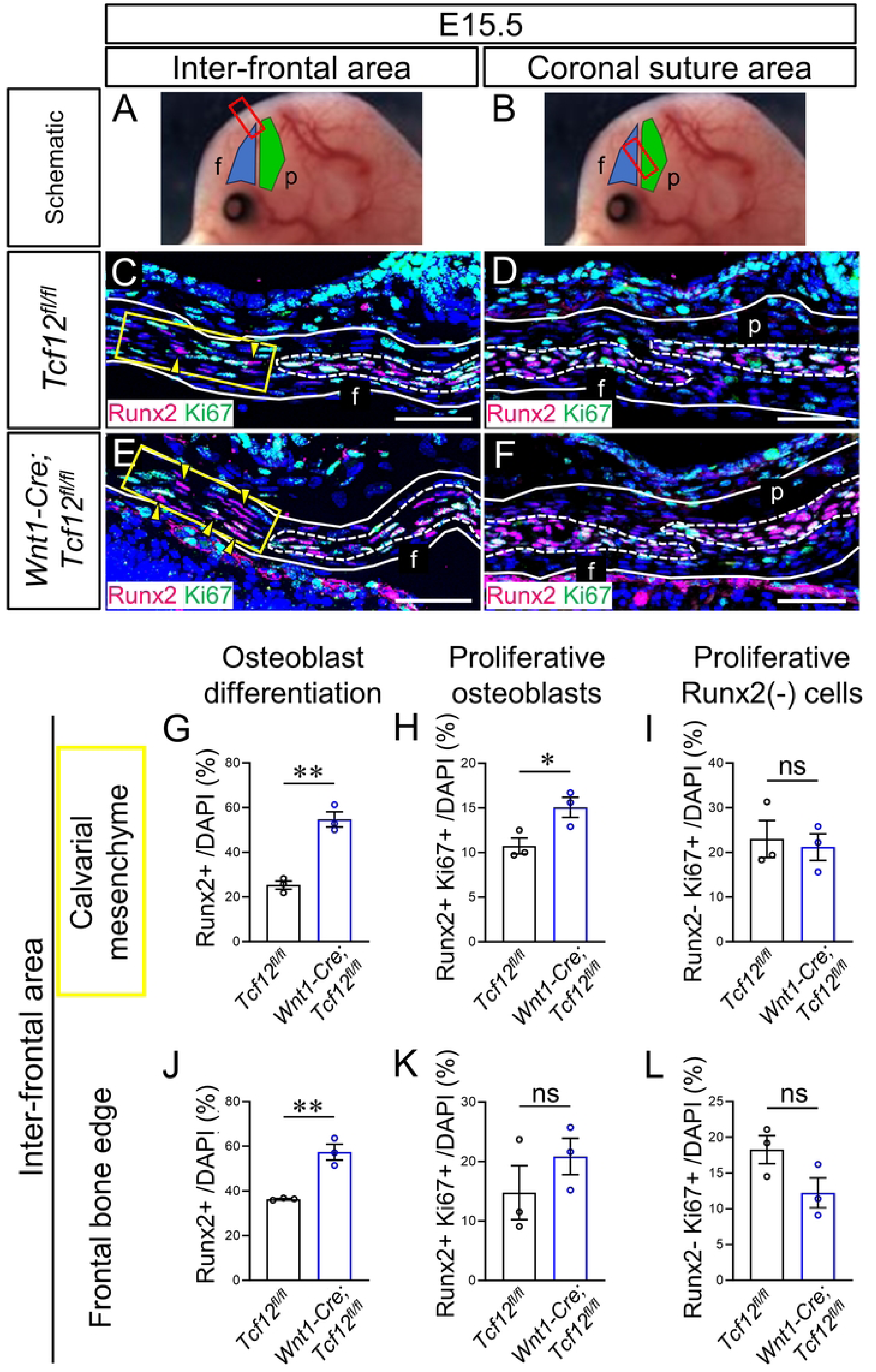
Loss of *Tcf12* in NCCs leads to upregulated osteogenic differentiation in the calvarial mesenchyme. (A, B) Schematic of an E15.5 mouse head with the frontal (blue) and parietal (green) bone rudiments. Red box shows the area of section for staining. (C-F) Runx2 (magenta) and Ki67 (green) immunofluorescence in the inter-frontal area (C, E) and the coronal suture area (D, F) of *Tcf12^fl/fl^* and *Wnt1-Cre;Tcf12^fl/fl^* mouse embryos at E15.5. Calvarial mesenchyme is outlined by white lines in C-F. Frontal bones and parietal bones are outlined by white dashed lines in C-F. Yellow arrowheads point to signals of Runx2 in C and E. (G-L) Quantification of the Runx2+ cells (G, J), Runx2+;Ki67+ cells (H, K), and Runx2-;Ki67+ cells (I, L) in the calvarial mesenchyme area at the level of posterior end of the inter-frontal suture of *Tcf12^fl/fl^* (n=3) and *Wnt1-Cre;Tcf12^fl/fl^* (n=3) mouse embryos at E15.5. Runx2+ / DAPI (%) is the percentage of Runx2+ cells out of all DAPI+ cells in the calvarial mesenchyme within 100 µm from the frontal bone edge to the apical side (yellow box) or to the basal side in C and E. Statistical differences were assessed with unpaired two-tailed Student’s t-test (G-L); ** = *P*-value <.01; * = <.05; ns, not significant. Error bars represent s.e.m. f, frontal bone; p, parietal bone. Scale bar = 50 μm.

### Osteogenic inhibitor *Lmx1b* expression is downregulated in the calvarial mesenchyme of *Wnt1-Cre;Tcf12^fl/fl^* mice

The ectopic bone formed in the apical part of the calvaria in *Wnt1-Cre;Tcf12^fl/fl^* mice was induced by upregulated osteogenic differentiation in the calvarial mesenchyme at embryonic stages. This phenotype was quite similar to one previously documented in mice with deletion of *Lmx1b* using *Prrx1-Cre*, which is widely expressed in the cranial mesenchyme; this deletion led to heterotopic ossification at the vertex as well as fusion of multiple sutures [20]. Furthermore, deletion of *Lmx1b* in NCCs, using *Sox10-Cre*, resulted in heterotopic bone and posterior extension of the frontal bone within the calvarial mesenchyme derived from NCCs [23]. *Lmx1b* is specifically expressed in the early migrating mesenchyme apical to the supra-orbital mesenchyme and serves to inhibit osteogenesis in that area [19,20]. To evaluate changes in the transcriptome and downstream targets of *Tcf12* following its deletion in the calvarial mesenchyme, we performed RNA sequencing (RNAseq) analysis of *Tcf12^fl/fl^* and *Wnt1-Cre;Tcf12^fl/fl^* mouse embryos at E12.5 (Fig 6A-6E). Due to their osteogenic differentiation defects, we focused on molecular markers associated with osteogenic fate determination. In the RNAseq results, we found that *Lmx1b* expression was significantly downregulated in the calvarial mesenchyme of *Wnt1-Cre;Tcf12^fl/fl^* mouse embryos at E12.5, while the osteogenic regulator *Twist1* and the osteogenic markers *Runx2* were not affected (Fig 6B-6E). Thus, we hypothesized that deletion of *Tcf12* leads to reduction of *Lmx1b* expression and then later upregulation of *Runx2* in the calvarial mesenchyme. To test this hypothesis, we focused on the expression patterns of *Lmx1b* in the calvarial mesenchyme in *Wnt1-Cre;Tcf12^fl/fl^* mice at E13.5 and E15.5. *Lmx1b* was widely expressed in the calvarial mesenchyme at E13.5 in control mice (Fig 6F), then the expression was limited to the apical part of the calvarial mesenchyme at E15.5 (Fig 6G). *Lmx1b* expression was comparatively reduced in the calvarial mesenchyme in *Wnt1-Cre;Tcf12^fl/fl^* mice at both E13.5 and E15.5 (Fig 6H and 6I). Additionally, *Tcf12* was co-expressed with *Lmx1b* in the calvarial mesenchyme of control mice, while the co-expression level of *Lmx1b* was decreased in *Wnt1-Cre;Tcf12^fl/fl^* mice at E13.5 (Fig 6J and 6K). These results indicated that *Lmx1b*, which is a potential downstream target of Tcf12, maybe directly regulated by *Tcf12* in the calvarial mesenchyme during calvarial bone development.

**Fig 6.**
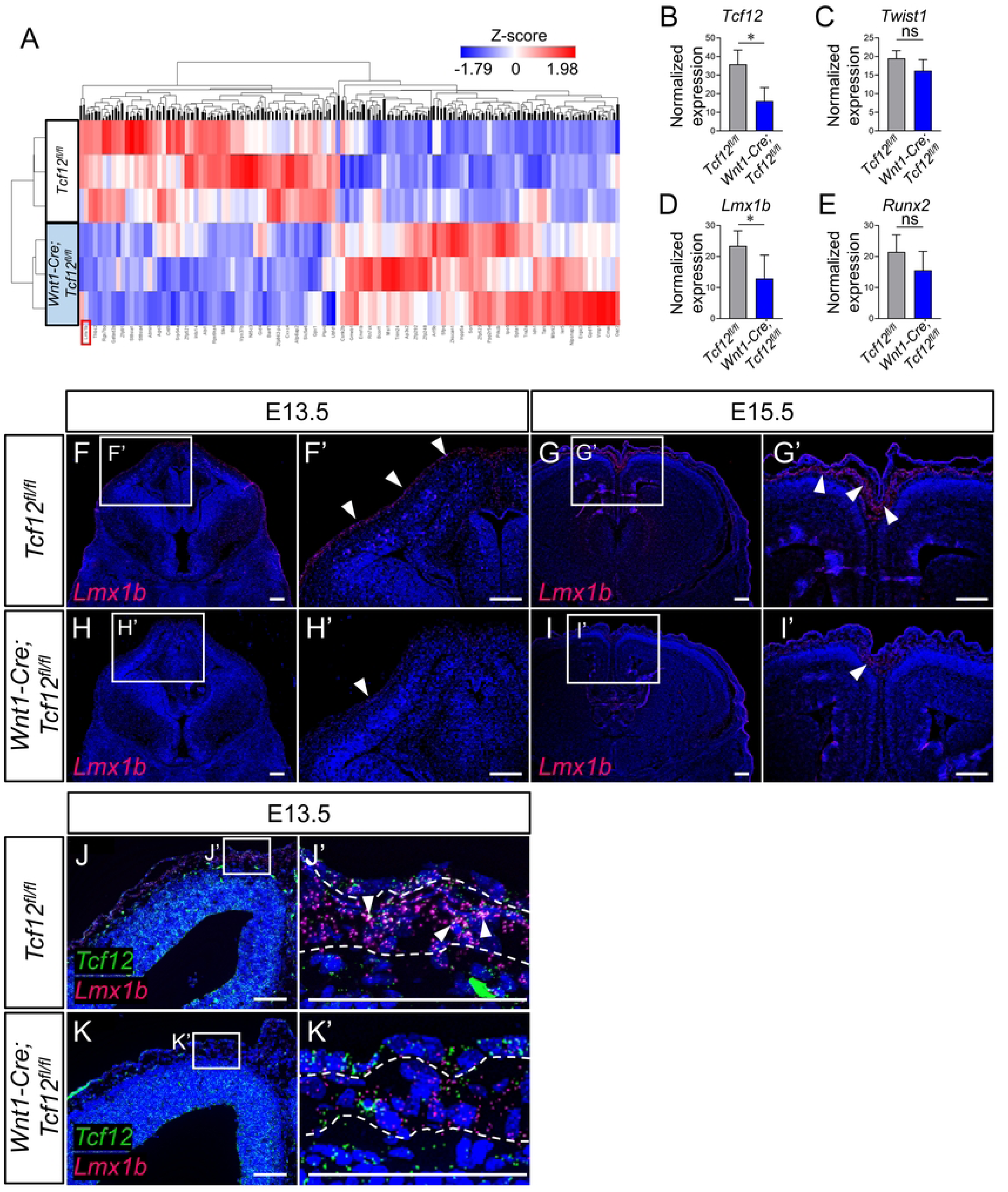
Loss of *Tcf12* in NCCs decreases *Lmx1b* expression levels in the calvarial mesenchyme. (A-E) RNAseq analysis of apical calvarial mesenchyme samples from *Tcf12^fl/fl^* (blue, upper 3 rows) and *Wnt1-Cre;Tcf12^fl/fl^* mice (red, lower 3 rows) collected at E12.5 with heat map (A) and normalized expression of *Tcf12* (B), *Twist1* (C), *Lmx1b* (D), and *Runx2* (E). Red rectangle highlights *Lmx1b* in A. Statistical differences were assessed with the algorithm in PartekFlow (B-E); * = *P* ≤ .05; ns, not significant. Error bars represent s.e.m. (F-I) *Lmx1b* expression patterns in the calvarial mesenchyme of *Tcf12^fl/fl^* and *Wnt1-Cre;Tcf12^fl/fl^* mouse embryos at E13.5 (F, H) and E15.5 (G, I). Highly magnified images within white boxes in F-I are indicated in F’-I’, respectively. White arrowheads point to the *Lmx1b* expression in F’-I’. (J, K) *Lmx1b* (magenta) and *Tcf12* (green) co-expression patterns in the calvarial mesenchyme of *Tcf12^fl/fl^* (J) and *Wnt1-Cre;Tcf12^fl/fl^* (K) mouse embryos at E13.5. Highly magnified images within white boxes in J and K are indicated in J’ and K’, respectively. White arrowheads point to the co-expression of *Lmx1b* and *Tcf12* in J’ and K’. Calvarial mesenchyme areas are outlined by white dashed lines in K’ and L’. Scale bars = 200 μm (F-I, F’-I’), 100 μm (J, K, J’, K’).

### Overexpression of *Lmx1b* reduces bone mineralization in the calvarial mesenchyme of *Wnt1-Cre;Tcf12^fl/fl^* mice

To investigate whether *Lmx1b* acts downstream of *Tcf12* in the regulation of calvarial bone formation, we conducted a rescue assay *in vitro*. We dissected the calvarial mesenchyme, excluding calvarial bones, from *Tcf12^fl/fl^* and *Wnt1-Cre;Tcf12^fl/fl^* mouse embryos at E13.5, then cultured the harvested cells. To achieve overexpression of *Lmx1b* in each group, plasmids with Lmx1b clone insertion and control vehicle were transfected. The *Lmx1b* expression level was efficiently increased in *Wnt1-Cre;Tcf12^fl/fl^* mouse embryos (Fig 7A). Using these cells, we conducted an osteogenic differentiation assay. Alizarin red staining after induction showed that overexpression of *Lmx1b* reduced the mineralization ability of cells from *Wnt1-Cre;Tcf12^fl/fl^* mouse embryos at E13.5 (Fig 7B-7E). Quantification of the alizarin red intensity showed that the intensity level was restored to a level comparable to that of controls in *Wnt1-Cre;Tcf12^fl/fl^* mouse embryos with *Lmx1b* overexpression (Fig 7F). To reveal how *Lmx1b* expression is controlled by Tcf12, we searched its promoter region for potential binding sites of Tcf12. Bioinformatic analysis using transcription binding profiles from the UCSC/JASPAR database showed that Tcf12 can potentially bind to the *Lmx1b* promoter region (Fig 7G). We utilized a CUT and RUN (Cleavage Under Targets and Release Using Nuclease) assay [24] and confirmed direct binding of Tcf12 to the predicted region at the *Lmx1b* promoter site (Fig 7H). Our data suggested that *Lmx1b* is a direct target of Tcf12 in the calvarial mesenchyme during calvarial bone development. Taken together, our findings indicated that *Lmx1b* acts downstream of Tcf12 to inhibit osteogenic differentiation of the calvarial mesenchymal cells that are present on the apical side of the frontal bone edge in early calvarial bone development.

**Fig 7.**
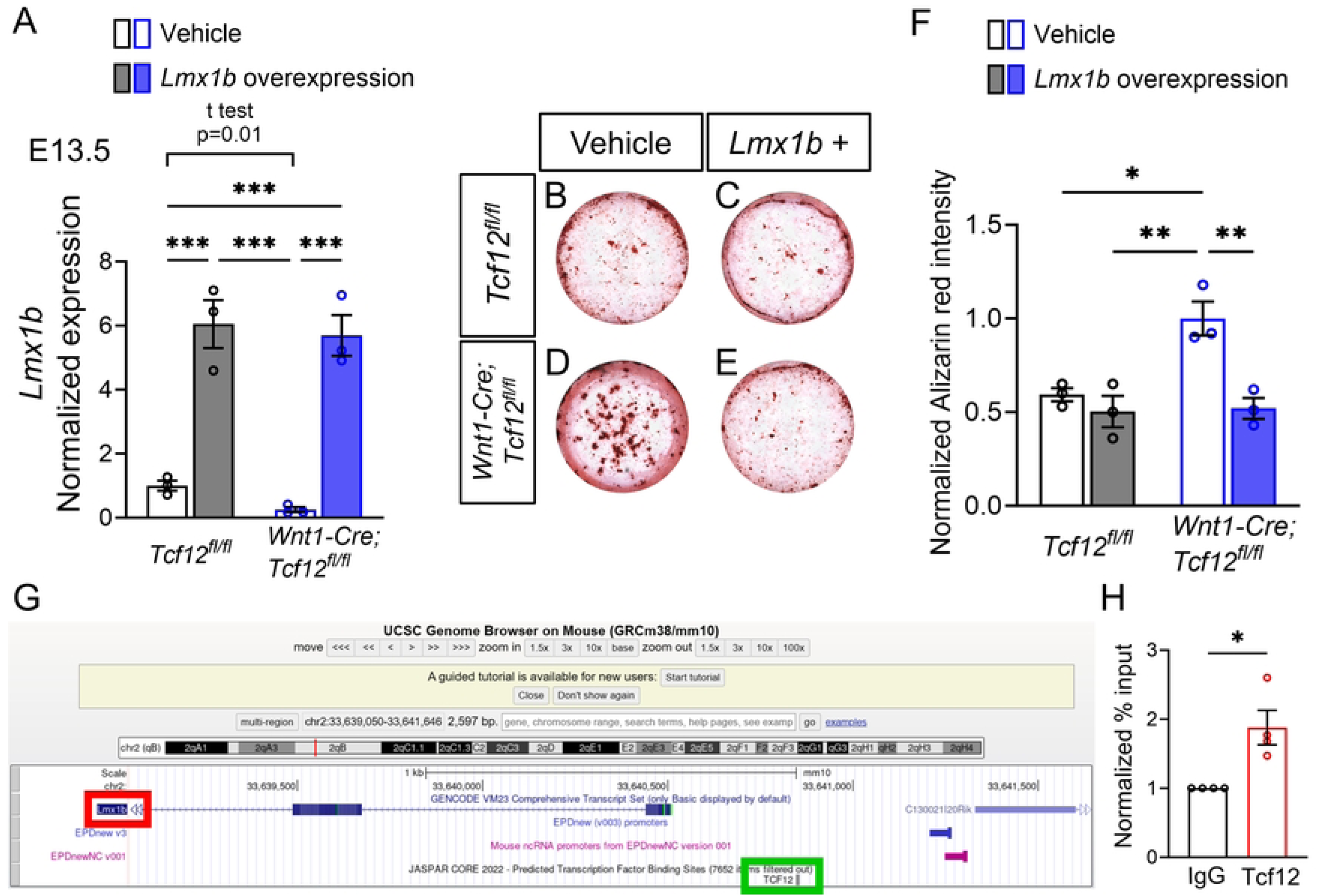
Overexpression of *Lmx1b* in the calvarial mesenchyme reduces osteogenic differentiation in *Wnt1-Cre;Tcf12^fl/fl^* cells *in vitro*. (A) qPCR analysis of *Lmx1b* expression in the calvarial mesenchyme of *Tcf12^fl/fl^* (black, n=3) and *Wnt1-Cre;Tcf12^fl/fl^* (blue, n=3) mouse embryos at E13.5 with transfection of vehicle (unfilled) and *Lmx1b* overexpression plasmid (filled) to confirm the efficiency of the overexpression. Values of *Lmx1b* expression are normalized with *Gapdh* and *Tcf12^fl/fl^* with vehicle transfection. (B-F) Alizarin Red S staining of cultured calvarial mesenchymal cells from *Tcf12^fl/fl^* (B, C) and *Wnt1-Cre;Tcf12^fl/fl^* (D, E) mouse embryos with transfection of vehicle and *Lmx1b* overexpression plasmid at E13.5 after osteogenic differentiation induction. Quantification of the alizarin red intensity in *Tcf12^fl/fl^* (n=3) and *Wnt1-Cre;Tcf12^fl/fl^* (n=3) mouse embryos at E13.5 with transfection of vehicle (unfilled) and *Lmx1b* overexpression plasmid (filled) (F). Values of alizarin red intensity are normalized with *Wnt1-Cre;Tcf12^fl/fl^* with vehicle transfection. (G, H) UCSC binding prediction of Tcf12 binding motif to *Lmx1b* (G). Red rectangle marks *Lmx1b* and green rectangle highlights Tcf12. CUT and RUN assay for binding of Tcf12 to *Lmx1b* promoter region in the calvarial mesenchyme of wild-type mouse embryos at E13.5 (n=4). Statistical differences were assessed with one-way ANOVA (A, F) or unpaired two-tailed Student’s t-test (*Tcf12^fl/fl^* with vehicle vs *Wnt1-Cre;Tcf12^fl/fl^* with vehicle in A, H); *** = *P*-value <.001; ** = <.01; * = <.05. Error bars represent s.e.m.

## Discussion

*TWIST1* and *TCF12* play important roles in regulating suture mesenchymal cells and suture patency in humans. A recent study has shown that haploinsufficiency of *Twist1* leads to craniosynostosis with neurocognitive deficits in mice [13]. Homozygous loss of *Tcf12* produces a similar calvarial phenotype to heterozygous loss of *Twist1*, with coronal suture fusion [12]. However, to date there has been no comprehensive analysis of the neurocognitive functions of *Tcf12* knockout models. In this study, we show impaired motor learning ability independent of craniosynostosis in *Tcf12* conditional knockout mouse models. Furthermore, we have investigated the frontal bone defects of these mice and found that *Tcf12* regulates osteogenesis in the calvarial mesenchyme through controlling *Lmx1b* expression during calvarial development.

There is partial coronal suture fusion which does not affect skull shape in *Wnt1-Cre;Mesp1-Cre;Tcf12^fl/fl^* mice at P1M. On the other hand, mice with heterozygous loss of *Twist1*, which encodes a binding partner of Tcf12, exhibit coronal suture fusion, and it is severe enough to affect skull shape [13,25–27]. In a recent study, we have shown that *Twist1^+/-^* mice with craniosynostosis have neurocognitive abnormalities, which are also seen in human Saethre-Chotzen syndrome [13]. In this study, we have discovered impaired motor learning ability and reduced cerebellar volume in mice with deletion of *Tcf12* in *Wnt1*-positive cells. The phenotype in these mice is limited to the cerebellum, where *Wnt1*-positive cells are present, while abnormalities in *Twist1^+/-^* mice with craniosynostosis are seen in multiple regions in the brain [13]. Notably, *Twist1* is not expressed in the mouse brain [13], whereas *Tcf12* is expressed in the cortex and cerebellum during development [28]. These findings suggest that the brain phenotypes in *Twist1^+/-^* and *Wnt1-Cre;Tcf12^fl/fl^* mice are due to different factors: craniosynostosis causing high intracranial pressure against the brain in the former and abnormal development of the brain in the latter. The pathogenesis behind abnormal development of the cerebellum in *Wnt1-Cre;Tcf12^fl/fl^* mice can be explained by the reduction of cell proliferation in the cerebellum, where there is specific deletion of *Tcf12*. However, further detailed analyses of the differentiation of the progenitor cells of the cerebellum and other aspects of cerebellar development will be necessary to reveal the mechanism of reduced cerebellar volume. Interestingly, intellectual disability, developmental delay, and learning disability have been reported previously in individuals with *TCF12* mutations regardless of whether they exhibit craniosynostosis [29]. It will be crucial to examine the expression pattern of TCF12 in the human brain and compare it to mice. Further functional studies on how TCF12 may be involved in controlling brain functions will be necessary to gain a better understanding of how *TCF12* mutation can lead to deficits in brain function. It should be noted that because our animal model did not target brain structures other than the cerebellum, loss of *Tcf12* in other regions could result in abnormal brain functions that would not be apparent in our mice.

In addition to coronal suture fusion in *Wnt1-Cre;Mesp1-Cre;Tcf12^fl/fl^* mice, there is posterior expansion of the frontal bones and ectopic bone formation in the medial part of the coronal suture area in *Wnt1-Cre;Tcf12^fl/fl^* mice at the adult stage, consistent with the phenotypes previously seen at younger stages [12]. Previous study has shown that haploinsufficiency of *Twist1* in the NCCs (*Wnt1-Cre;Twist1^fl/+^* mice) also leads to frontal bone expansion [30]. These similarities between *Tcf12* and *Twist1* mutant mice imply that *Tcf12* and *Twist1* may play similar roles or act together in frontal bone and coronal suture development. Expanded frontal bone or ectopic bone with inverted overlap between the frontal and parietal bones in *Wnt1-Cre;Tcf12^fl/fl^* mice are present approximately half of the sagittal suture and the apical coronal suture, which have mesenchyme derived from NCCs [23,31]. This suggests that not only the frontal bone primordia but also perhaps the suture mesenchyme or calvarial mesenchyme derived from NCCs are involved in frontal bone expansion and ectopic bone formation in *Wnt1-Cre;Tcf12^fl/fl^* mice. At the cellular and molecular level, we have revealed that frontal bone expansion in *Wnt1-Cre;Tcf12^fl/fl^* mice is caused by the acceleration of frontal bone growth with excess osteoblast production in the frontal bone edge [12], as well as by an ectopic osteogenic center (marked by Runx2+ cells) in the direction of frontal bone expansion within the calvarial mesenchyme derived from NCCs. Notably, ectopic bone seen in *Wnt1-Cre;Tcf12^fl/fl^* mouse embryos is derived from NCCs, from which the frontal bones also originate [12]. These findings indicate that the deletion of *Tcf12* promotes the differentiation of the mesenchymal stem cells into osteoblasts in the apical calvarial mesenchyme. The ectopic bone may fuse to the forming frontal bones, eventually forming posteriorly expanded frontal bones.

In the calvarial mesenchyme during embryonic development, *Lmx1b* is specifically expressed in the early migrating mesenchyme apical to the supra-orbital mesenchyme and inhibits osteogenesis in this region [19,20]. Deletion of *Lmx1b* in the entire calvarial mesenchyme leads to ectopic osteogenesis in the early migrating mesenchyme, and deletion of *Lmx1b* in NCCs results in heterotopic bone and posterior extension of the frontal bone within the calvarial mesenchyme derived from NCCs [20,23], similar to what we have seen in *Wnt1-Cre;Tcf12^fl/fl^* mice. We show that the expression of *Lmx1b*, which is a potential downstream target of Tcf12, is reduced in the calvarial mesenchyme in *Wnt1-Cre;Tcf12^fl/fl^* mouse embryos at E12.5, E13.5 and E15.5, then osteogenic differentiation marker *Runx2* is upregulated by E15.5. Cell proliferation, including that of all cell populations in the early migrating mesenchyme, is not affected in *Lmx1b* knockout mice [20]. Here we showed that the total number of proliferating cells is not affected in this area in *Wnt1-Cre;Tcf12^fl/fl^* mice, but the proliferation of Runx2+ cells increases, suggesting that *Tcf12* might negatively regulate proliferation of early osteogenic progenitor cells expressing *Runx2* in the apical part of the calvarial mesenchyme at the embryonic stage as well as inhibiting osteogenic differentiation in this region. Overexpression of *Lmx1b* in *Tcf12* mutant calvarial mesenchymal cells led to normal levels of osteogenic differentiation, whereas untreated *Tcf12* mutant cells exhibited overactive osteogenic differentiation caused by reduced *Lmx1b* expression. This is consistent with a previous study which has shown that overexpression of *Lmx1b* in osteoblast precursor cells from P1 mouse calvaria inhibits osteogenic differentiation through the BMP2 signaling pathway; conversely, the knockdown of *Lmx1b* in these cells enhances osteogenic differentiation [32]. We have also shown that osteogenic differentiation of the calvarial mesenchymal cells on the apical side of the frontal bone edge may be negatively regulated by *Lmx1b* via direct binding of Tcf12 to the *Lmx1b* promoter site in early calvarial bone development in mice. Alongside this role, *Lmx1b* is necessary for the lineage commitment (apical-basal patterning) of the cranial mesenchyme at E12.5, and acts downstream of *Tcf12* in the regulation of the apical expansion of the calvarial bones after this stage. Although previous study has shown that silencing *Tcf12* promotes osteoblastic differentiation, upregulating *Runx2* expression levels in bone mesenchymal stem cells via BMP and Erk1/2 signaling pathways [33], our study links Tcf12 directly with Lmx1b in controlling osteogenic differentiation during calvarial bone morphogenesis.

In summary, our study points to roles of *Tcf12* in controlling calvarial bone formation and cerebellar development in mice, partially recapitulating phenotypes of patients with *TCF12* mutation, including impaired brain function even in the absence of craniosynostosis. These results represent important findings that eventually will lead to more effective treatment options and more accurate prognoses concerning neurocognitive dysfunctions after surgery for craniosynostosis patients with impaired brain function. Moreover, our study has revealed a novel regulatory mechanism through which *Tcf12* and *Lmx1b* control osteogenic differentiation in the apical part of the calvarial mesenchyme where the future frontal and parietal bones expand. Future experiments will be required to identify more detailed molecular and cellular mechanisms underlying proper coronal suture formation and calvarial bone expansion toward the apex.

## Materials and Methods

### Animals

Mouse experiments were approved by the University of Southern California Institutional Animal Care and Use Committee and performed following the regulations for animal experiments (Protocol 21329). All mice were housed under a 12 h light/dark cycle in pathogen-free conditions with free access to food and water, in accordance with the Guide for Care and Use of Laboratory Animals of the National Institutes of Health. All the mice were identified by ear tags. Ear biopsies were lysed at 55℃ overnight in DirectPCR solution (Viagen, 102-T) followed by 85℃ heat inactivation for 1 hour and PCR-based genotyping (GoTaq Green Master Mix, Promega, and C1000 Touch Cycler, Bio-rad). *Tcf12* [34], *Wnt1-Cre* [35], *Mesp1-Cre* [36] and alleles were genotyped by PCR. Wild-type and mutant mice were maintained on a C57BL/6J background. Mice were euthanized by carbon dioxide overdose and then decapitation. For all experiments, mice of both sexes were used.

### Behavioral assays

Behavioral tests were carried out at 8-12 weeks of age, using protocols as previously described [13] with minor modifications. In all experiments, mice were acclimated to the behavior room at least 60 min before the first trial began. For each assay, 10 males and 10 females were used. Experimenters were blinded to animal genotypes during behavioral tests and data analyses.

### Rotarod test

The rotarod test consists of training and test phases. Mice were first trained by placing them on a rotating rod (Panlab, Harvard Apparatus) at a constant speed of 4 rpm until they were able to stay on the rotating rod for 60 s. The test phase was performed 24 hours after the training phase. The rotarod apparatus was set to accelerate from 4 to 40 rpm in 300 s, and mice were placed on the rod initially rotating at 4 rpm. The latency (time) to falling off the rod was determined. Each mouse was tested three times a day at 15 min intervals for four consecutive days.

### Novel object test

This test was performed as described previously [37] with minor modifications. Briefly, this test consists of habituation, familiarization, and test phases. In the habituation phase, subject mice were placed in the center of a clean, empty cage and allowed to explore freely for 10 min. After 24 h, the familiarization phase was performed. Two identical objects were taped to the floor along the long axis, 10 cm from the south and north walls. The mouse was placed in the center of the cage facing the east or west wall and allowed to explore for 10 min. The test phase was performed 24 h after the familiarization phase. One of the identical objects was replaced with a novel object with a different shape but similar size. The mouse was placed in the center of the cage facing the east or west wall and allowed to explore for 10 min. The apparatus and objects were thoroughly cleaned with 70% ethanol. The entire test phase was videotaped and the travel of the subject mouse was manually documented. The preference index was calculated as (T_n_ – T_f_)/(T_n_ + Tf) × 100%, where T_n_ and T_f_ represent the time spent exploring novel and familiar objects, respectively.

### Y maze test

This test was performed as described previously [38] with minor modifications. Briefly, the Y maze apparatus (Panlab, Harvard Apparatus) consisted of three closed arms (30 × 6 cm). Mice were individually placed in the A arm, facing the center, and allowed to freely explore the apparatus for 5 min. The apparatus and objects were thoroughly cleaned with 70% ethanol. A, B, and C arm entries (all four paws in an arm) were manually recorded. The spontaneous alteration, indicating the mouse’s willingness to explore, was calculated as [The number of times the mouse entered different arms 3 times in a row]/([Total entries] – 2) × 100%.

### Elevated plus maze test

This test was performed as described previously [39,40] with minor modifications. Briefly, the elevated plus maze apparatus (Panlab, Harvard Apparatus) consisted of two open arms and two closed arms (29.5 × 6 cm). The entire maze was elevated 40 cm from the floor. Mice were individually placed in the center of the maze, facing the open arm, and allowed to freely explore the apparatus for 10 min. The apparatus and objects were thoroughly cleaned with 70% ethanol. Open and closed arm entries (all four paws in an arm) and the time spent in the open arms were recorded using Smart v3.0 (Panlab, Harvard Apparatus). The open arm entry score was calculated as (Open entries)/(Total entries) × 100%.

### Three-chamber social interaction test

The test apparatus was a Plexiglas box containing three compartments connected by small openings that allow the mice free access to each compartment. The subject mouse was first placed in the middle chamber and allowed to explore the three empty chambers for 10 min. After this habituation, the mouse was gently guided to the middle chamber and side doors were closed. A stranger mouse was placed in an inverted wire cup in one side chamber and an empty wire cup was placed in the other side chamber. Then the side doors were opened and the subject mouse was allowed to freely explore the chambers for 10 min. After this period, the subject mouse was again guided to the middle chamber and the doors were closed. A second stranger mouse was placed in the previously empty wire cup. The side doors were opened, and the subject mouse was allowed to freely explore for another 10 min. The apparatus and wire cups were thoroughly cleaned with 70% ethanol. The entire test phase was videotaped and the travel of the subject mouse was manually documented. The amount of time that the subject mouse spent sniffing each wire cup was quantified and the preference index was calculated as (T_s1_ – T_e_)/(T_s1_ + T_e_) × 100% and (T_s2_ – T_s1_)/(T_s2_ + T_s1_) × 100%. T_e_, T_s1_ and T_s2_ represent the time spent exploring the empty, Stranger 1 and Stranger 2 wire cups, respectively.

### Micro-CT (µCT) analysis

µCT imaging and Generalized Procrustes analysis of postnatal skulls were performed using identical conditions as previously described [13] with minor modifications.

The calvarial bones were radiographed in one-month-old live mice using a micro-CT (Ct Lab In-vivo 90) device. Images were collected using a 70 KVp and 114 mA X-ray source. All the 3D reconstructions and sections were analyzed using AVIZO 9.4.0 (Thermo Fisher Scientific).

Generalized Procrustes analysis was conducted to reveal the spatial configuration of the samples using MorphoJ 1.07a [41] referencing previous work [25]. The groups compared were *Tcf12^fl/fl^* mice, *Wnt1-Cre;Tcf12^fl/fl^* mice, *Mesp1-Cre;Tcf12^fl/fl^* mice, *Wnt1-Cre;Mesp1-Cre;Tcf12^fl/fl^* mice (without coronal suture fusion), and *Wnt1-Cre;Mesp1-Cre;Tcf12^fl/fl^* mice (with coronal suture fusion). Subsets of landmarks were used to individually analyze the differences in shape of two distinct regions of the skull (top of calvarium and lateral portion of calvarium). Principal component analysis was used to identify the variation between different groups in both these distinct regions. The landmark subsets are indicated in S1 Fig.

### Magnetic resonance imaging (MRI) analysis

MRI and volume measurement of brains of three-month-old mice were conducted as previously described [13]. All mice were anesthetized with 4% isoflurane and intracardially perfused with 30 mL of 0.1 M PBS containing 10 U/ml heparin (Sigma, H3149) and 2 mM ProHance (a Gadolinium contrast agent, Bracco Diagnostics, 0270-1111-01), followed by 30 mL of 4% paraformaldehyde (PFA) containing 2 mM ProHance. Then the mice were decapitated, and brains with skulls were incubated in 4% PFA + 2 mM ProHance overnight at 4℃, then transferred to 0.1 M PBS containing 2 mM ProHance and 0.02% sodium azide for 10 days before MRI scanning [42]. MRI images were acquired on an MRSolutions 7 Tesla MRI scanner (Guildford, UK). Three-dimensional anatomical Fast Spin Echo (FSE) images were acquired for the whole brain. Imaging parameters were as follows: TEeffective/TR = 26ms/400ms, 4 averages, echo train length = 4, field of view = 16 mm × 16 mm × 25.6 mm, matrix size = 160 × 256 × 256, and flip angle = 90°. Total acquisition time was 280 minutes. The brain region-of-interest boundaries were manually drawn for each slice using ImageJ. The measured areas of the whole brain, cerebellum, isocortex, and hippocampus were multiplied by slice thickness to calculate final volumes (mm^3^).

### Histology, Immunohistochemistry (IHC) staining and *in situ* **hybridization (ISH)**

Suture samples were dissected and fixed in 10% neutralized buffered formalin (Sigma-Aldrich, HT501128) overnight at room temperature, then decalcified with 10% EDTA in PBS for about 2 weeks. Decalcified calvaria were dehydrated with graded EtOH solutions (from 70% to 100%) followed by xylene, then embedded in paraffin. Tissue blocks were sectioned at 7 μm using a microtome (Leica) and mounted on SuperFrost Plus slides (Fisher, 12-550-15). Hematoxylin and Eosin (H&E) staining was performed following standard protocols.

Samples were fixed with 4% paraformaldehyde (PFA) in PBS overnight at 4℃, followed by dehydration with graded sucrose solutions (15% and 30% sucrose in PBS, each overnight at 4℃) and immediately embedded in O.C.T. compound (Sakura Finetek, 4583). For adult suture samples, decalcification in 10% EDTA in PBS was performed for about 2 weeks after fixation. Frozen tissue blocks were sectioned at 8 mm on a cryostat (Leica) and mounted on SuperFrost Plus slides (VWR, 48311-703) for staining.

For IHC, sections were treated with sodium citrate buffer (Vector, H-3300-250) at 98°C for 5 min and permeabilized with blocking buffer containing 1% BSA, 2% goat serum and 0.3% Triton X-100 in PBS for 1 h at room temperature, then incubated overnight at 4℃ with the following primary antibodies: anti-Runx2 (Santa Cruz Biotechnology SC390715, 1:50), anti-Sp7 (Abcam ab22552, 1:100), or anti-Ki67 (Abcam ab15580, 1:200). The following day, sections were incubated for 2 h at room temperature with fluorescently conjugated secondary antibodies: Alexa Fluor 488 goat anti-rabbit IgG (Invitrogen A11008, 1:200) or Alexa Fluor 647 goat anti-mouse IgG (Invitrogen A21235, 1:200). Counterstaining was then conducted using DAPI (Thermo Fisher Scientific 62248, 1:1000), followed by mounting with Immu-Mount (Fisher Scientific, 9990402).

ISH was assayed using RNAScope Multiplex Fluorescent Detection Kit v2 (Advanced Cell Diagnostics, 323110). Mm-Tcf12 (504861) and Mm-Lmx1b (412931) probes from Advanced Cell Diagnostics were used in this study. Sections were acquired as mentioned above, after which ISH was performed according to the manufacturer’s instructions.

Images were captured by a Leica DMI3000 B microscope and a Keyence BZ-X810 system.

### Alizarin Red S staining of P0 skulls

Skulls of P0 mice were stained for bone with 2% Alizarin Red S in 1% KOH for about 3 days. The specimens were then cleared in a glycerol series containing 1% KOH and stored in 100% glycerol. Samples were imaged using a Leica M125 microscope.

### RNA sequencing analysis

The calvarial mesenchyme, excluding supra-orbital mesenchyme, from E12.5 embryos was dissected. The ectoderm and the brain were carefully removed manually. Tissues were kept in RNALater (Thermo Fisher Scientific, AM7020) at −80°C. Tissue from 3 to 4 embryos was pooled to make one sample. RNeasy Micro Kit (Qiagen, 74004) was used to isolate RNA. UCLA Technology Center for Genomics and Bioinformatics performed the RNA quality tests, cDNA library preparation and sequencing (Illumina Novaseq SP, 2 × 50 bp read length). Differential expression was filtered by selecting transcripts with fold change ≥ |1.5| and a significance level of P ≤ 0.05 using PartekFlow.

### Calvarial mesenchyme primary cell culture

The calvarial mesenchyme, excluding supra-orbital mesenchyme, from E13.5 embryos was dissected. One embryo was used for each replicate. To isolate the mesenchyme, the surface ectoderm and the brain were removed carefully. The mesenchyme was then washed with PBS and αMEM with 100 U/ml penicillin and 100 µg/ml streptomycin several times, minced into tiny pieces and transferred into a T25 dish (Nest, 705001) at 37℃ in an atmosphere of 5% CO_2_. The cell culture medium contained αMEM (Gibco, 12571063) supplemented with 20% FBS (Biowest, S1620), 100 U/ml penicillin and 100 µg/ml streptomycin (Gibco, 15140122). After 7 days, cells were digested with TrypLE (Gibco, 12604013) and the tissues were removed by passing through a 70 µm cell strainer (Falcon, 352350), then the cells were incubated for each experiment.

To induce overexpression of *Lmx1b* in the cells, Lmx1b Mouse Tagged ORF Clone plasmids (ORIGENE, MG226016) were transfected using Lipofectamine LTX Reagent with PLUS Reagent (Invitrogen, 15338100) according to the manufacturer’s protocol. We used pCMV6-AC-GFP Mammalian Expression Vector (ORIGENE, PS100010) as control vehicle plasmids.

### Quantitative real-time PCR (qPCR)

Total RNA was extracted from the cells after transfection using the RNeasy Micro Kit (Qiagen, 74004). RNA was reverse transcribed using an iScript cDNA Synthesis Kit (Bio-Rad, 1708890). Quantitative real-time PCR was performed using SsoFast EvaGreen Supermixes (Bio-Rad, 1725200). Primers for qPCR of *Lmx1b* were the sequences F: 5’-TTCCTGATGCGAGTCAACGAG-3’ and R: 5’-TCCGATCCCGGAAGTAGCAG-3’. *Gapdh* was used as a reference gene for normalization. Gene expression was quantified with the ΔCT method.

### Osteogenic differentiation assay

The cells were seeded into 48-well plates with αMEM containing 10% FBS, 100 U/ml penicillin and 100 µg/ml streptomycin. Osteogenic differentiation was promoted by incubation in osteogenic medium containing αMEM supplemented with 2.5% FBS, 10 nM Dexamethasone, 10 mM β-glycerophosphate, and 50 µg/ml Ascorbic acid for about 7 days. The culture medium was replaced every 2 days. For Alizarin red staining, cells were fixed with 100% MeOH, then stained with 1% Alizarin Red S solution. Samples were captured using a Keyence BZ-X810 system.

### UCSC/JASPAR binding prediction and Cut and Run assay

Predicted sites of Tcf12 binding to the promoter region of the *Lmx1b* gene were identified at chr2: 33640847-33640857, using the UCSC/JASPAR bioinformatics database. The primers used for the Cut and Run assay were forward 5’-CTACCGGCTACTCGCAGC-3’ and reverse 5’-CACTGGAGTAGTGCGGGGA-3’. The cells were collected according to the manufacturer’s instructions for the Cut and Run assay kit (Cell Signaling, 86652). 10 μl of IgG isotype control (Cell Signaling, 3900) or Tcf12 (HEB) (Santa Cruz Biotechnology, sc-365980) were used for each reaction. One embryo was used for each replicate.

### Statistics

GraphPad Prism Software (Prism 9) was used for statistical analysis. Each data point in the bar graphs represents one biological replicate. For each qPCR experiment, at least two technical replicates were analyzed. For histological staining, immunofluorescence, and ISH, we used samples from at least three individual mice/embryos, and representative images are shown. All bar graphs display mean ± s.e.m. Statistical comparisons were done using unpaired two-tailed Student’s t-test for comparison of two groups or one-way ANOVA for comparison of more than two groups. P ≤ 0.05 and < 0.05 were treated as statistically significant for the RNAseq analysis and all other analyses, respectively.

## Acknowledgements

The authors thank Bridget Samuels for critical reading of the manuscript and Drs. Robert Maxson and Pedro Sanchez for their helpful discussions.

## Supporting information

**S1 Fig. Landmarks for analyzing head shape (Related to** Fig 2). (A-C) Landmarks for analyzing head shape. Table of landmark descriptions for analyzing head shape (A). Superior (B) and lateral (C) views of landmark locations.

**S2 Fig. Loss of *Tcf12* in *Wnt1-*expressing cells has no effect outside the cerebellum in mice.** (A-E) No significant differences were seen in the novel object test (A), Y maze test (B), elevated plus maze test (C), or three-chamber test (D, E) in comparisons of *Tcf12^fl/fl^* (black), *Wnt1-Cre;Tcf12^fl/fl^* (blue), *Mesp1-Cre;Tcf12^fl/fl^* (green), and *Wnt1-Cre;Mesp1-Cre;Tcf12^fl/fl^* (red) mice at P2M-P3M. Each group, n=20 including 10 males and 10 females. (F, G) Quantification of the brain volume acquired by MRI images (in Fig. 3C-F) of the cortex (F) and the hippocampus (G) in *Tcf12^fl/fl^* (n=3), *Wnt1-Cre;Tcf12^fl/fl^* (n=3), *Mesp1-Cre;Tcf12^fl/fl^* (n=3), and *Wnt1-Cre;Mesp1-Cre;Tcf12^fl/fl^* (n=3) mice at P3M. (H-K) Ki67 (green) immunofluorescence on the cortex and the hippocampus in *Tcf12^fl/fl^* (H) and *Wnt1-Cre;Tcf12^fl/fl^* (I) mouse embryos at E18.5. Quantification of the Ki67+ cells in the cortex (J) and the hippocampus (K) of *Tcf12^fl/fl^* (n=3) and *Wnt1-Cre;Tcf12^fl/fl^* (n=3) mouse embryos at E18.5. Ki67+ cell (%) is the percentage of Ki67 positive cells out of all cells in the yellow box on the cortex in J and in the red circle on the hippocampus in K. (L, M) *Tcf12* expression patterns in the developing brain in *Tcf12^fl/fl^* (L) and *Wnt1-Cre;Tcf12^fl/fl^* (M) mouse embryos at E18.5. Panels at top right in L and M show higher magnifications of the white dotted rectangle areas. Yellow arrowheads point to the *Tcf12* expression in L and M. Statistical differences were assessed with one-way ANOVA (A-G) or unpaired two-tailed Student’s t-test (J, K); ns, not significant. Error bars represent s.e.m. cx, cortex; hip, hippocampus; ce, cerebellum. Scale bar = 200 μm (H, I, L, M).

**S3 Fig. *Tcf12* expression in the apical part of the calvarial mesenchyme is efficiently deleted in *Wnt1-Cre;Tcf12^fl/fl^* mouse embryos at E15.5.** (A) Schematic of an E15.5 mouse head with the frontal (blue) and parietal (green) bone rudiments. Red box shows the area of section for staining. (B, C) *Tcf12* expression pattern in the posterior portion of the inter-frontal suture mesenchyme in *Tcf12^fl/fl^* (B) and *Wnt1-Cre;Tcf12^fl/fl^* (C) mouse embryos at E15.5. Calvarial mesenchyme areas are outlined by white dashed lines in B and C. Asterisk in C indicates the deleted expression of *Tcf12* in the calvarial mesenchyme. f, frontal bone; p, parietal bone. Scale bar = 100 μm.

**S4 Fig. Loss of *Tcf12* in NCCs does not affect osteogenic differentiation or cell proliferation in the frontal and parietal bone edges of the coronal suture in mouse embryos.** (A-F) Quantification of the Runx2+ cells (A, D), Runx2+;Ki67+ cells (B, E), and Runx2-;Ki67+ cells (C, F) in the coronal suture area of *Tcf12^fl/fl^* (n=3) and *Wnt1-Cre;Tcf12^fl/fl^* (n=3) mouse embryos at E15.5. Runx2+ / DAPI (%) is the percentage of Runx2+ cells out of all DAPI+ cells within 100 µm on the frontal bone edge in Figure 5D and F. Statistical differences were assessed with unpaired two-tailed Student’s t-test; ns, not significant. Error bars represent s.e.m.

## References

1. Ishii M, Sun J, Ting MC, Maxson RE. The Development of the Calvarial Bones and Sutures and the Pathophysiology of Craniosynostosis. Curr Top Dev Biol. 2015;115:131–56. doi: 10.1016/bs.ctdb.2015.07.004. PMID: 26589924.

2. Twigg SR, Wilkie AO. A Genetic-Pathophysiological Framework for Craniosynostosis. Am J Hum Genet. 2015 Sep 3;97(3):359–77. doi: 10.1016/j.ajhg.2015.07.006. PMID: 26340332.

3. Stanton E, Urata M, Chen JF, Chai Y. The clinical manifestations, molecular mechanisms and treatment of craniosynostosis. Dis Model Mech. 2022 Apr 1;15(4):dmm049390. doi: 10.1242/dmm.049390. PMID: 35451466.

4. Gripp KW, Zackai EH, Stolle CA. Mutations in the human TWIST gene. Hum Mutat. 2000;15(2):150–5. doi: 10.1002/(SICI)1098-1004(200002)15:2<150::AID-HUMU3>3.0.CO;2-D. Erratum in: Hum Mutat 2000;15(5):479. PMID: 10649491.

5. Morriss-Kay GM, Wilkie AO. Growth of the normal skull vault and its alteration in craniosynostosis: insights from human genetics and experimental studies. J Anat. 2005 Nov;207(5):637–53. doi: 10.1111/j.1469-7580.2005.00475.x. PMID: 16313397.

6. Brooks ED, Beckett JS, Yang J, Timberlake AT, Sun AH, Chuang C, et al. The Etiology of Neuronal Development in Craniosynostosis: A Working Hypothesis. J Craniofac Surg. 2018 Jan;29(1):49–55. doi: 10.1097/SCS.0000000000004040. PMID: 29049144.

7. Howard TD, Paznekas WA, Green ED, Chiang LC, Ma N, Ortiz de Luna RI, et al. Mutations in TWIST, a basic helix-loop-helix transcription factor, in Saethre-Chotzen syndrome. Nat Genet. 1997 Jan;15(1):36–41. doi: 10.1038/ng0197-36. PMID: 8988166.

8. el Ghouzzi V, Le Merrer M, Perrin-Schmitt F, Lajeunie E, Benit P, Renier D, et al. Mutations of the TWIST gene in the Saethre-Chotzen syndrome. Nat Genet. 1997 Jan;15(1):42–6. doi: 10.1038/ng0197-42. PMID: 8988167.

9. Sharma VP, Fenwick AL, Brockop MS, McGowan SJ, Goos JA, Hoogeboom AJ, et al. Mutations in TCF12, encoding a basic helix-loop-helix partner of TWIST1, are a frequent cause of coronal craniosynostosis. Nat Genet. 2013 Mar;45(3):304–7. doi: 10.1038/ng.2531. Erratum in: Nat Genet. 2013 Oct;45(10):1261. PMID: 23354436; PMCID: PMC3647333.

10. Fan X, Waardenberg AJ, Demuth M, Osteil P, Sun JQJ, Loebel DAF, et al. TWIST1 Homodimers and Heterodimers Orchestrate Lineage-Specific Differentiation. Mol Cell Biol. 2020 May 14;40(11):e00663–19. doi: 10.1128/MCB.00663-19. PMID: 32179550.

11. Connerney J, Andreeva V, Leshem Y, Muentener C, Mercado MA, Spicer DB. Twist1 dimer selection regulates cranial suture patterning and fusion. Dev Dyn. 2006 May;235(5):1345–57. doi: 10.1002/dvdy.20717. Erratum in: Dev Dyn. 2012 Feb;241(2):433. PMID: 16502419.

12. Ting MC, Farmer DT, Teng CS, He J, Chai Y, Crump JG, et al. Embryonic requirements for Tcf12 in the development of the mouse coronal suture. Development. 2022 Jan 1;149(1):dev199575. doi: 10.1242/dev.199575. PMID: 34878091.

13. Yu M, Ma L, Yuan Y, Ye X, Montagne A, He J, et al. Cranial Suture Regeneration Mitigates Skull and Neurocognitive Defects in Craniosynostosis. Cell. 2021 Jan 7;184(1):243–256.e18. doi: 10.1016/j.cell.2020.11.037. PMID: 33417861.

14. Ferguson JW, Atit RP. A tale of two cities: The genetic mechanisms governing calvarial bone development. Genesis. 2019 Jan;57(1):e23248. doi: 10.1002/dvg.23248. PMID: 30155972.

15. Jiang X, Iseki S, Maxson RE, Sucov HM, Morriss-Kay GM. Tissue origins and interactions in the mammalian skull vault. Dev Biol. 2002 Jan 1;241(1):106–16. doi: 10.1006/dbio.2001.0487. PMID: 11784098.

16. Yoshida T, Vivatbutsiri P, Morriss-Kay G, Saga Y, Iseki S. Cell lineage in mammalian craniofacial mesenchyme. Mech Dev. 2008 Sep-Oct;125(9-10):797–808. doi: 10.1016/j.mod.2008.06.007. PMID: 18617001.

17. Sasaki T, Ito Y, Bringas P Jr, Chou S, Urata MM, Slavkin H, et al. TGF-beta-mediated FGF signaling is crucial for regulating cranial neural crest cell proliferation during frontal bone development. Development. 2006 Jan;133(2):371–81. doi: 10.1242/dev.02200. PMID: 16368934.

18. Deckelbaum RA, Holmes G, Zhao Z, Tong C, Basilico C, Loomis CA. Regulation of cranial morphogenesis and cell fate at the neural crest-mesoderm boundary by engrailed 1. Development. 2012 Apr;139(7):1346–58. doi: 10.1242/dev.076729. PMID: 22395741.

19. Roybal PG, Wu NL, Sun J, Ting MC, Schafer CA, Maxson RE. Inactivation of Msx1 and Msx2 in neural crest reveals an unexpected role in suppressing heterotopic bone formation in the head. Dev Biol. 2010 Jul 1;343(1-2):28–39. doi: 10.1016/j.ydbio.2010.04.007. PMID: 20398647.

20. Cesario JM, Landin Malt A, Chung JU, Khairallah MP, Dasgupta K, Asam K, et al. Anti-osteogenic function of a LIM-homeodomain transcription factor LMX1B is essential to early patterning of the calvaria. Dev Biol. 2018 Nov 15;443(2):103–116. doi: 10.1016/j.ydbio.2018.05.022. PMID: 29852132.

21. Dasgupta K, Chung JU, Asam K, Jeong J. Molecular patterning of the embryonic cranial mesenchyme revealed by genome-wide transcriptional profiling. Dev Biol. 2019 Nov 15;455(2):434–448. doi: 10.1016/j.ydbio.2019.07.015. PMID: 31351040.

22. Zhuang Y, Cheng P, Weintraub H. B-lymphocyte development is regulated by the combined dosage of three basic helix-loop-helix genes, E2A, E2-2, and HEB. Mol Cell Biol. 1996 Jun;16(6):2898–905. doi: 10.1128/MCB.16.6.2898. PMID: 8649400.

23. Cabrera Pereira A, Dasgupta K, Ho TV, Pacheco-Vergara M, Kim J, Kataria N, et al. Lineage-specific mutation of *Lmx1b* provides new insights into distinct regulation of suture development in different areas of the calvaria. Front Physiol. 2023 Aug 1;14:1225118. doi: 10.3389/fphys.2023.1225118. PMID: 37593235.

24. Skene PJ, Henikoff S. An efficient targeted nuclease strategy for high-resolution mapping of DNA binding sites. Elife. 2017 Jan 16;6:e21856. doi: 10.7554/eLife.21856. PMID: 28079019.

25. Parsons TE, Weinberg SM, Khaksarfard K, Howie RN, Elsalanty M, Yu JC, et al. Craniofacial shape variation in *Twist1^+/-^* mutant mice. Anat Rec (Hoboken). 2014 May;297(5):826–33. doi: 10.1002/ar.22899. PMID: 24585549.

26. Nuri T, Ota M, Ueda K, Iseki S. Quantitative Morphologic Analysis of Cranial Vault in *Twist1^+/-^* Mice: Implications in Craniosynostosis. Plast Reconstr Surg. 2022 Jan 1;149(1):28e–37e. doi: 10.1097/PRS.0000000000008665. PMID: 34936613.

27. Hoshino Y, Takechi M, Moazen M, Steacy M, Koyabu D, Furutera T, et al. Synchondrosis fusion contributes to the progression of postnatal craniofacial dysmorphology in syndromic craniosynostosis. J Anat. 2023 Mar;242(3):387–401. doi: 10.1111/joa.13790. PMID: 36394990.

28. Uittenbogaard M, Chiaramello A. Expression of the bHLH transcription factor Tcf12 (ME1) gene is linked to the expansion of precursor cell populations during neurogenesis. Brain Res Gene Expr Patterns. 2002 Jan;1(2):115–21. doi: 10.1016/s1567-133x(01)00022-9. PMID: 15018808.

29. Davis EE, Balasubramanian R, Kupchinsky ZA, Keefe DL, Plummer L, Khan K, et al. TCF12 haploinsufficiency causes autosomal dominant Kallmann syndrome and reveals network-level interactions between causal loci. Hum Mol Genet. 2020 Aug 11;29(14):2435–2450. doi: 10.1093/hmg/ddaa120. PMID: 32620954.

30. Teng CS, Ting MC, Farmer DT, Brockop M, Maxson RE, Crump JG. Altered bone growth dynamics prefigure craniosynostosis in a zebrafish model of Saethre-Chotzen syndrome. Elife. 2018 Oct 25;7:e37024. doi: 10.7554/eLife.37024. PMID: 30375332.

31. Doro D, Liu A, Grigoriadis AE, Liu KJ. The Osteogenic Potential of the Neural Crest Lineage May Contribute to Craniosynostosis. Mol Syndromol. 2019 Feb;10(1-2):48–57. doi: 10.1159/000493106. PMID: 30976279.

32. Kim K, Kim JH, Kim I, Seong S, Han JE, Lee KB, et al. Transcription Factor Lmx1b Negatively Regulates Osteoblast Differentiation and Bone Formation. Int J Mol Sci. 2022 May 7;23(9):5225. doi: 10.3390/ijms23095225. PMID: 35563615.

33. Yi S, Yu M, Yang S, Miron RJ, Zhang Y. Tcf12, A Member of Basic Helix-Loop-Helix Transcription Factors, Mediates Bone Marrow Mesenchymal Stem Cell Osteogenic Differentiation In Vitro and In Vivo. Stem Cells. 2017 Feb;35(2):386–397. doi: 10.1002/stem.2491. PMID: 27574032.

34. Wojciechowski J, Lai A, Kondo M, Zhuang Y. E2A and HEB are required to block thymocyte proliferation prior to pre-TCR expression. J Immunol. 2007 May 1;178(9):5717–26. doi: 10.4049/jimmunol.178.9.5717. PMID: 17442955.

35. Danielian PS, Muccino D, Rowitch DH, Michael SK, McMahon AP. Modification of gene activity in mouse embryos in utero by a tamoxifen-inducible form of Cre recombinase. Curr Biol. 1998 Dec 3;8(24):1323–6. doi: 10.1016/s0960-9822(07)00562-3. PMID: 9843687.

36. Saga Y, Miyagawa-Tomita S, Takagi A, Kitajima S, Miyazaki Ji, Inoue T. MesP1 is expressed in the heart precursor cells and required for the formation of a single heart tube. Development. 1999 Aug;126(15):3437–47. doi: 10.1242/dev.126.15.3437. PMID: 10393122.

37. Leger M, Quiedeville A, Bouet V, Haelewyn B, Boulouard M, Schumann-Bard P, et al. Object recognition test in mice. Nat Protoc. 2013 Dec;8(12):2531–7. doi: 10.1038/nprot.2013.155. PMID: 24263092.

38. Kraeuter AK, Guest PC, Sarnyai Z. The Y-Maze for Assessment of Spatial Working and Reference Memory in Mice. Methods Mol Biol. 2019;1916:105–111. doi: 10.1007/978-1-4939-8994-2_10. PMID: 30535688.

39. Holmes A, Wrenn CC, Harris AP, Thayer KE, Crawley JN. Behavioral profiles of inbred strains on novel olfactory, spatial and emotional tests for reference memory in mice. Genes Brain Behav. 2002 Jan;1(1):55–69. doi: 10.1046/j.1601-1848.2001.00005.x. PMID: 12886950.

40. Holmes A, Yang RJ, Crawley JN. Evaluation of an anxiety-related phenotype in galanin overexpressing transgenic mice. J Mol Neurosci. 2002 Feb-Apr;18(1-2):151–65. doi: 10.1385/JMN:18:1-2:151. PMID: 11931346.

41. Klingenberg CP. MorphoJ: an integrated software package for geometric morphometrics. Mol Ecol Resour. 2011 Mar;11(2):353–7. doi: 10.1111/j.1755-0998.2010.02924.x. PMID: 21429143.

42. Gompers AL, Su-Feher L, Ellegood J, Copping NA, Riyadh MA, Stradleigh TW, et al. Germline Chd8 haploinsufficiency alters brain development in mouse. Nat Neurosci. 2017 Aug;20(8):1062–1073. doi: 10.1038/nn.4592. PMID: 28671691.

